# Interpretable Chirality-Aware Graph Neural Network for Quantitative Structure Activity Relationship Modeling in Drug Discovery

**DOI:** 10.1101/2022.08.24.505155

**Authors:** Yunchao “Lance” Liu, Yu Wang, Oanh Vu, Rocco Moretti, Bobby Bodenheimer, Jens Meiler, Tyler Derr

**Affiliations:** Vanderbilt University; Leipzig University

## Abstract

In computer-aided drug discovery, quantitative structure activity relation models are trained to predict biological activity from chemical structure. Despite the recent success of applying graph neural network to this task, important chemical information such as molecular chirality is ignored. To fill this crucial gap, we propose <u>Mol</u>ecular-<u>K</u>ernel <u>G</u>raph <u>N</u>eural <u>N</u>etwork (MolKGNN) for molecular representation learning, which features SE(3)-/conformation invariance, chirality-awareness, and interpretability. For our MolKGNN, we first design a molecular graph convolution to capture the chemical pattern by comparing the atom’s similarity with the learnable molecular kernels. Furthermore, we propagate the similarity score to capture the higher-order chemical pattern. To assess the method, we conduct a comprehensive evaluation with nine well-curated datasets spanning numerous important drug targets that feature realistic high class imbalance and it demonstrates the superiority of MolKGNN over other GNNs in CADD. Meanwhile, the learned kernels identify patterns that agree with domain knowledge, confirming the pragmatic interpretability of this approach. Our codes are publicly available at https://github.com/meilerlab/MolKGNN.

## 1 Introduction

Developing new drugs is time-consuming and expensive, e.g., it took cabozantinib, an oncologic drug, 8.8 years and $1.9 billion to get on the market (Prasad and Mailankody 2017). To assist this process, computer-aided drug discovery (CADD) has been widely used. In CADD, several mathematical and machine learning methods have been developed to model the Quantitative Structure Activity Relationship (QSAR) to predict the biological activity of molecules based on their geometric structures (Sliwoski et al. 2014).

Recently, Graph Neural Networks (GNNs) have successfully been applied in many fields, e.g., social networks and recommender systems (Zhou et al. 2020). As molecules can be essentially viewed as graphs with atoms as nodes and chemical bonds as edges, GNNs are naturally adopted to perform such graph classification, i.e., predicting biological activity of molecules based on their geometric structures (Atz, Grisoni, and Schneider 2021). A typical GNN architecture for graph classification begins with an encoder extracting node representations by passing neighborhood information followed by pooling operations that integrate node representations into graph representations, which are fed into a classifier to predict graph classes (Zhou et al. 2020).

Despite the promise of GNN models applied to molecular representation learning, existing GNN models either blindly follow the message passing framework without considering molecular constraints on graphs (Feng et al. 2022), fail to integrate chirality (Schütt et al. 2017), or lack interpretability (Adams, Pattanaik, and Coley 2021). To fill this crucial gap, we develop a new GNN model named MolKGNN that features SE(3)/conformation invariance, chirality-awareness and provides a form of interpretability. Our main contributions can be summarized as follows:

- **Novel Interpretable Molecular Convolution:** We design a new convolution operation to capture chemical pattern of each atom by quantifying the similarity between the atom’s neighboring subgraph and the learnable molecular kernel, which is inherently interpretable.
- **Better Chirality Characterization:** Rather than listing all permutations of neighbors for a chiral center, or using dihedral angles, the chirality calculation module in our design only needs a lightweight linear algebra calculation.
- **Comprehensive Evaluation in CADD:** We perform a comprehensive evaluation using well-curated datasets spanning numerous important drug targets (that feature realistic high class imbalance) and metrics that bias predicted active molecules for actual experimental validation. Ultimately, we demonstrate the superiority of MolKGNN over other GNNs in CADD.

## 2 Related Work

### 2.1 Extending Convolutions to the Graph Domain

Convolutional Neural Networks (CNN) have enjoyed much success on images. However, convolution fails to readily extend to graphs due to their irregular structures. Early efforts on GNNs focused on spectral convolution (Defferrard, Bresson, and Vandergheynst 2016; Kipf and Welling 2016). Later, spatial-based methods define graph convolution based on nodes’ spatial relationship (Gilmer et al. 2017).

Vanilla GNN is known to have limited expressive power bounded by the Weisfeiler-Lehman (WL) graph isomorphism test (Xu et al. 2018) and hence have difficulty in finding substructures. On the other hand, graph kernels can take substructures into consideration by computing a similarity score among graph substructures. Recently, a strand of work extend GNNs by combining them with graph kernels to distinguish substructures (Cosmo et al. 2021; Feng et al. 2022). However, we argue that extending the expressive power to distinguish more substructures is not necessarily helpful with molecular representation learning. For example, (Cosmo et al. 2021) explicitly states their model could distinguish triangles. Nevertheless, although present in some drug molecules (Talele 2016), triangles are rare due to chemical instability. This can be verified by an empirical observation in the annual best-selling small molecule drugs posters^1^ (McGrath, Brichacek, and Njardarson 2010). Moreover, learning a useful discrete structure in a differentiable way is challenging and hence the structure learning process in (Cosmo et al. 2021) uses random modification. This raises the question of whether it is worth the extra computational time associated with finding a structural similarity. Instead, our method identifies semantic similarity between a 1-hop molecular neighborhood and a molecule kernel (Figure. 3).

**Figure 1:**
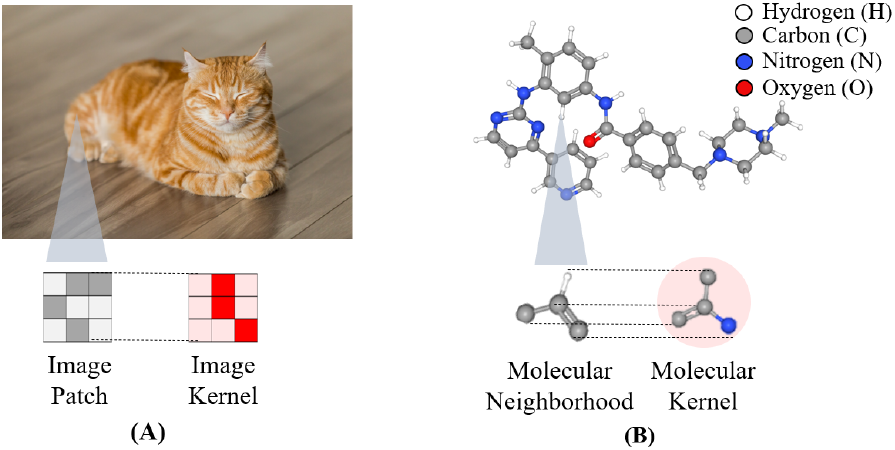
Analogy between (A) 2D image convolution and (B) 3D molecular convolution. In (A), the more similar the image patch is to the image kernel, the higher the output value. We design molecular convolution (B) to output a higher value if the molecular neighborhood is similar to the molecular kernel. The kernel provides the basis for our chirality calculation and has added benefit of interpretability.

**Figure 2:**
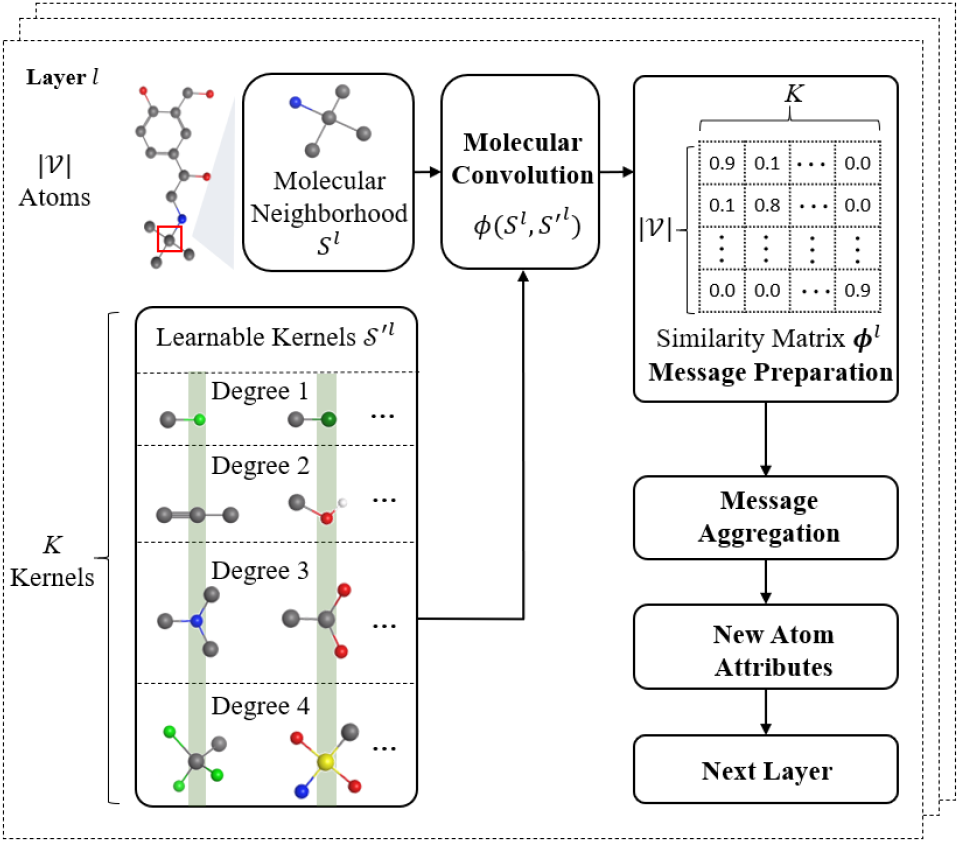
An overview of the proposed MolKGNN. For each atom 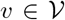 of the molecule *G*, its 1-hop star-like molecular neighborhood subgraph *S^l^* is convoluted with a set of *K* learnable kernels 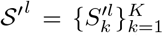 to get its similarity vector and collectively for all nodes, after applying the same convolution as above, we end up with the similarity matrix 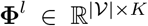 in layer *l*. Specifically, 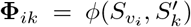 quantifies the similarity between the neighborhood subgraph around atom *υ_i_* and the *k*^th^ kernel (See Fig. 3 for more details of calculating 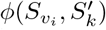). The **Φ**^*l*^ serves as the new atom attributes for the computation in the next layer *l* + 1.

**Figure 3:**
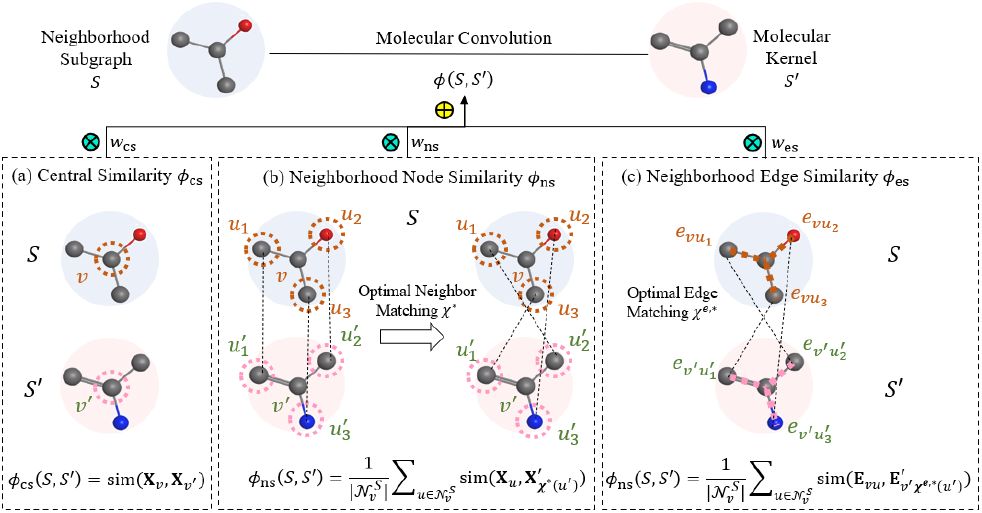
Illustration of the molecular convolution. The similarity between a neighborhood subgraph *S* and a kernel *S*^′^ is quantified by *ϕ*(*S, S*’). This similarity score is calculated as the combination of *ϕ*_cs_, *ϕ*_ns_, *ϕ*_es_, which quantify the similarity of center node, neighboring nodes, edges, respectively.

### 2.2 Molecular Representation Learning

It is not surprising to see the application of GNN to molecules due to the ready interpretation of atoms as nodes and bonds as edges. Even the term *graph* (in the sense used in graph theory) was used for the first time to draw a relationship between mathematics and chemistry (Sylvester 1878). Several attempts have been made to leverage GNNs for molecular representation learning. In this paper, we classify them into four categories.

Models in the first category capture the 2D connectivity (i.e., molecular constitution). Examples include (Yang et al. 2019; Coley et al. 2017). Some molecular properties, especially pharmacological activities, are dependent on the chirality of molecules (H Brooks, C Guida, and G Daniel 2011).

A chiral molecule cannot be superimposed on its own mirror image. For such tasks, molecules should be treated as noninvariant to reflection. Models in this second category are reflection-sensitive, or chirality-aware and sometimes called 2.5D QSAR models. Examples include (Liu et al. 2021; Pattanaik et al. 2020). However, molecules are not planar graphs but are 3D entities. Due to rotations around single bonds, molecules can display different conformations. Models in this third category, e.g., (Flam-Shepherd et al. 2021), take the dihedral angles of rotatable bonds into consideration to distinguish different conformations. As molecules exist as conformational ensembles, a fourth category (4D) encodes ensembles instead of individual conformations (Adams, Pattanaik, and Coley 2021).

## 3 Preliminaries and Problem Definition

In this section, we introduce all notations used throughout this paper and define the problem of QSAR modeling.

### Notations

We represent a molecule as an attributed and undirected graph 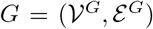 where 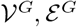 are the set of nodes (atoms) and edges (chemical bonds). Let 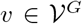 denote the node *υ* and 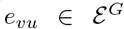 denote an edge between *υ* and *u*. Moreover, we represent the node attribute matrix as 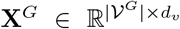 and edge attribute matrix as 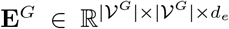 where *d_υ_,d_e_* are the dimension of node and edge features. Specifically, we let 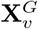 be the attribute of node *υ* and 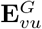 be the attribute of edge *e_υu_*. The node coordinate matrix is represented as 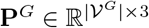 and 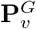 denotes the 3D coordinates of *υ*. The graph topology is described by its adjacency matrix 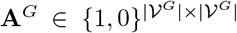 where 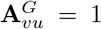 if 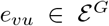, and 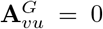 otherwise. Note that bond types are encoded as edge features. Furthermore, we denote the 1-hop neighborhood of *υ* in *G* as 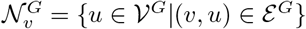.

### Problem Definition

Based on the above notations, we formulate QSAR modeling as a graph classification problem, which can be mathematically defined as: *Given a set of attributed molecule graphs* 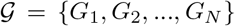 *with each molecule* 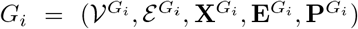 *as defined above and its corresponding one-hot encoded label* **Y**_*i*_, *we aim to learn a graph encoder* 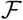 *and a classifier* 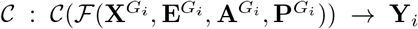 that is well-predictive of the ground truth label **Y**_*i*_ of molecule *G_i_*.

## 4 Molecular-Kernel Graph Neural Network

In this section, we introduce the framework of our proposed MolKGNN. As shown in Figure 2, MolKGNN recursively performs molecular convolution and message aggregation to learn representations of each molecule. In molecular convolution, we design learnable molecular kernels to capture chemically-meaningful subgraph pattern of each node/atom. Specifically, we calculate the similarity scores of each atom with its neighborhood to the molecular kernels and treat the obtained score as new atom features, which essentially describes the distance of the atom’s chemical properties to the patterns encoded in the kernels. Then in message aggregation, we leverage feature propagation to aggregate similarity scores of neighborhoods to further capture chemical context of each atom. These two modules proceed alternatively to gradually enlarge the receptive field so that we can capture higher-order chemical pattern. Next, we introduce details of molecular convolution and message aggregation.

### 4.1 Molecular Convolution

In 2D images, convolution operation can be regarded as calculating the similarity between the image patch and the image kernel. Larger output values indicate higher visual similarity patterns such as edges, strips, curves (Lin, Huang, and Wang 2021). Inspired by that, we design a molecular convolution that outputs higher values when a molecular neighborhood and kernels are more chemically similar (Figure 1). However, performing convolution on the irregular neighborhood subgraphs requires the learnable molecular kernels to have correspondingly different geometrical structures, which is computationally prohibitive. To handle this challenge, for each atom *υ* of degree *d* in *G*, we only consider its 1-hop star-like neighborhood subgraph 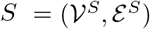 where 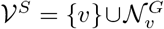 and 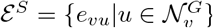. To make the molecular convolution feasible, we initialize the molecular kernel to also follow star-structure and denote it as 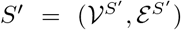 where 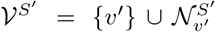 with *υ*’ being the central node without loss of generality and 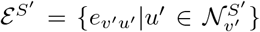. Let the learnable feature matrix and edge feature matrix of *S*’ be 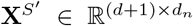 and 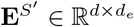, respectively. Then we define the operation of molecular convolution between the atom *υ* and the molecular kernel *S*^′^ as quantifying the similarity *ϕ* between *υ*’s neighborhood subgraph *S* and the kernel *S*′:

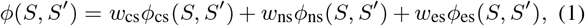

where *ϕ*_cs_,*ϕ*_ns_, *ϕ*_es_ quantify the similarity from three different aspects: the central similarity, neighborhood similarity, and edge similarity. We combine them together with learnable weights *w*_cs_,*w*_ns_,*w*_es_ ∈ [0,1] after softmax-normalization.

#### Central Similarity

We first capture the chemical property of the atom *υ* itself in *S* by computing its similarity to the central node *υ*′ in the kernel *S*′:

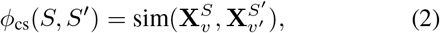

where 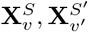 are attributes of the central atom *υ* in the subgraph S and the central node *υ*′ in the kernel *S*′. The sim(·, ·) is the function measuring vector similarity and we use cosine similarity throughout this work.

#### Neighboring Node and Edge Similarity

Besides the central node, the chemical property of an atom is also impacted by its neighborhood context, which motivates us to further quantify the similarity between 1) the neighboring nodes 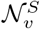 in *S* and 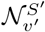 in *S*’, and 2) the edges 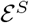 and 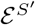.

Before calculating *φ*_ns_, *φ*_es_ between *S* and *S*′, we face a matching problem. For example, in Figure 3, the node *u*_1_ in *S* has more than one matching candidates, i.e., 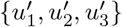 in *S*’. Here we seek a bijective matching 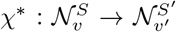 such that the average attribute similarity between 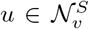 and 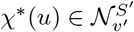 over all neighbors can be maximized:

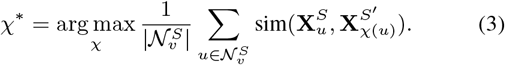

Given that exhausting all 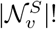 possible matchings to find the optimal one is computationally infeasible, we significantly simplify this computation by constraining the searching space according to the inherent structure of molecules, which are: 1) node degrees in drug-like molecule graphs are usually less than 5, with most atoms having a degree of 1 and few nodes having a degree of 4 (Patrick 2013); 2) for nodes of degree 4, only 12 among the total 24 possible matchings are valid after considering chirality (Pattanaik et al. 2020) (See Section 6.1 for more details).

After we obtain the optimal node matching, the bijective edge matching can be trivially defined as: 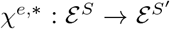 such that the edge 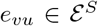 if and only if 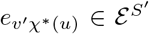. Then, with the defined node and edge matching *χ*\*,*χ*^*e*,*^ as above, we compute *ϕ*_ns_ and *ϕ*_es_ as:

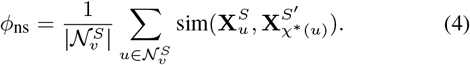

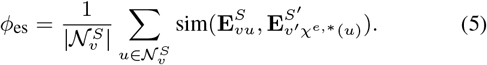

#### Chirality Characterization

After we capture the chemical-informative pattern of the neighborhood subgraph around each atom by quantifying the above three different aspects of similarity, our model is still chirality-insensitive, i.e., it still cannot distinguish enantiomers (pairs of mirror-imaged molecules that are non-superimposable, like our left and right hands (McNaught, Wilkinson et al. 1997)) However, chirality is a key determinant of a molecule’s biological activity (Sliwoski et al. 2012), which motivates us to characterize the chirality of the molecule in the next.

According to the rule of basic chemistry, chirality can only exist when the central atom has four unique neighboring substructures. Therefore, we only characterize the chirality for atom *ν* where 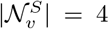 and its four neighboring substructures are different from each other. More specifically, given the neighborhood subgraph of an atom *S* forming the tetrahedron shown in Figure 4 where the four unique neighboring atoms are 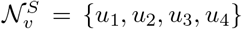, we select *u*_1_ without loss of generality as the anchor neighbor to define the three concurrent sides of the tetrahedron 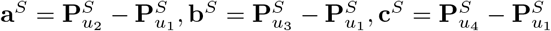 and further calculate the tetrahedral volume of *S* as:

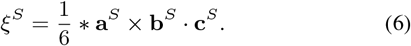

**Figure 4:**
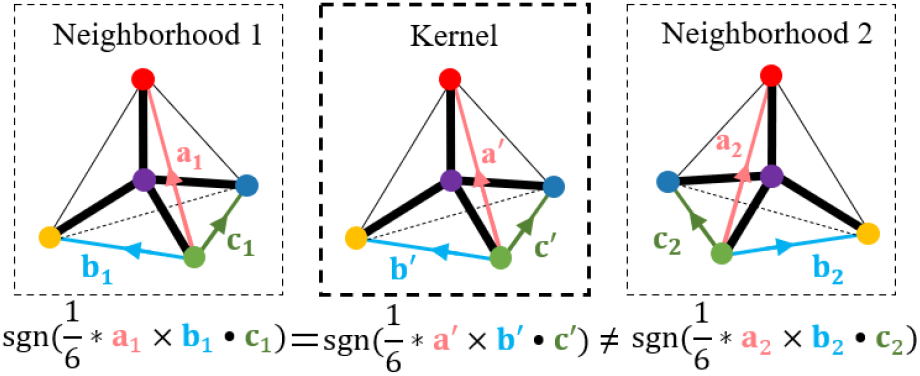
Illustration of chirality calculation. Corresponding nodes given by the optimal matching *χ*\* are of the same colors. sgn(·) is the sign function. We can distinguish mirror-imaged neighborhoods of two atoms by comparing the orientation of their corresponding tetrahedrons (i.e., the sign of the tetrahedral volume).

Similarly, we calculate 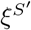 for the kernel *S*′. Since the sign of the tetrahedron volume of the molecule *ξ^S^* defines its orientation (Sliwoski et al. 2012), if sgn(*ξ^S^*) = -sgn(*ξ*^*S*′^), with sgn(·) being the sign function, the four neighboring subgraph *S, S*′ would be of opposite direction. Therefore, by treating the kernel as the anchor and comparing its direction with the ones of two neighborhood subgraphs as:

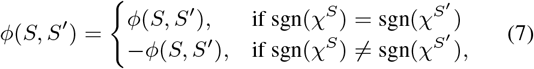

we can distinguish two enantiomers and make the model chirality-sensitive.

After we encode the chirality into the similarity computation, our proposed molecular convolution could fully capture the chemical pattern of the atom in terms of its own property by *ϕ*_cs_(*S, S*′), its neighborhood property *ϕ*_ns_(*S, S*′), *ϕ*_es_(*S, S*’) and its chirality by the sign of *ϕ*(*S, S*’). Since one kernel can only characterize one chemical pattern, we extend the above defined molecular convolution to the situation of multiple kernels in the next section.

### 4.2 Model Architecture

Suppose the set of *K* kernels at layer *l* be 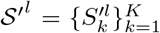, we apply the proposed molecular convolution with the learnable molecular kernel 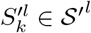 over the node representation **H**^*l*–1^ at the previous layer *l* –1 to obtain the node similarity matrix at layer *l* as 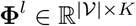:

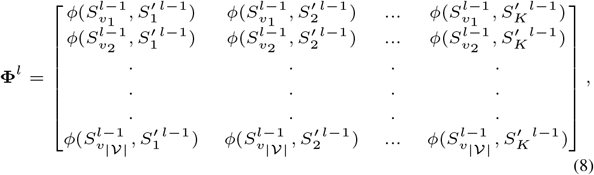

where 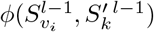 defines the similarity between the neighborhood subgraph around the atom *υ_i_* and the *k*^th^ kernel at layer *l* – 1. We note that 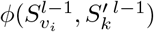 is set to 0 if 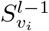 and 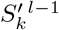 have different degrees so that back-propagation keeps the parameters in kernels of different degree untouched.

The above molecular kernel convolution can only capture the chemical pattern embedded in the 1-hop neighborhood around each atom. To further discover the chemical pattern embedded in the multi-hop neighborhood, we leverage the message-passing and directly aggregate the calculated neighborhood similarity **Φ**^*l*^ as:

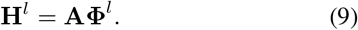

After recursively alternating between the molecular convolution and the message-passing *L* layers, the final atom rep-resentation **H**^*l*^ describes the chemical pattern up to *L* hops away of each atom. We further apply a readout function to integrate node presentations into the graph representation **G** for each graph *G* as:

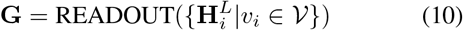

Here we employ global-sum pooling as our READOUT function, which adds all nodes’ representations.

### 4.3 Model Optimization

From now, let the graph representation of *G_i_* be **G**_*i*_ and so we apply the above process to get the representations for all *N* labeled graphs in 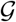. Then given the one-hot encoded label matrix **Y** ∈ ℝ^*N*×*C*^, the overall objective function of MolKGNN is formally defined as:

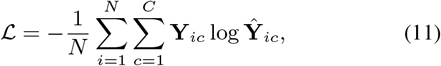

where 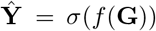 and *f* (·) is a classifier, e.g., Multi-Layer Perceptron followed by a softmax normalization *σ*.

## 5 Experiments

The experiments discussed in this section answer the following research questions:

- RQ1: How does MolKGNN compare against existing methods in a realistic drug discovery benchmark?
- RQ2: Do the learned kernels of MolKGNN align with domain knowledge and provide further interpretability?
- RQ3: Does MolKGNN possess the expressiveness for distinguishing chiral molecules?

### 5.1 A Realistic Drug Discovery Scenario

High-throughput screening (HTS) is the use of automated equipment to rapidly screen thousands to millions of molecules for the biological activity of interest in the early drug discovery process (Bajorath 2002). However, this brute-force approach has low hit rates, typically around 0.05%-0.5%. QSAR models are trained on the results of HTS experiments in order to screen additional molecules virtually and prioritize these for acquisition (Mueller et al. 2010; Butkiewicz et al. 2019). Dataset challenges for constructing QSAR models include imbalance (many more inactive molecules) (Wang et al. 2021) and containing false positives/negatives (Baell and Holloway 2010). Thus, for developing QSAR methods, curated high-quality datasets are needed. Unfortunately, too often small, uncurated, or unrealistic datasets are used, e.g., the commonly used ogbg-molpcba (Hu et al. 2020), which had no preprocessing to remove potential experimental artifacts (Ramsundar et al. 2015). Another commonly-used large molecule dataset is OGB-LSC PCQM4Mv2 (~ 3M molecules) (Hu et al. 2021), but is of molecular properties instead of biological activities. Lastly, when assessing the performance of QSAR models, a metric that bias the molecules with the highest predicted activities is of interest as ultimately only these will be acquired or synthesized and tested.

Fortunately, the CADD community has over the past 30 years developed suitable benchmark datasets and metrics that should be adopted in the machine learning community.

### 5.2 Datasets

PubChem (Kim et al. 2021) is a database supported by the National Institute of Health (NIH) that contains biological activities for millions of drug-like molecules, often from HTS experiments. However, the raw primary screening data from the PubChem have a high false positive rate (Butkiewicz et al. 2017, 2013). A series of secondary experimental screens on putative actives is used to remove these. While all relevant screens are linked, the datasets of molecules are not curated to list all inactive molecules from the primary HTS and only confirmed actives after secondary screening (Butkiewicz et al. 2017, 2013). Thus, we identified nine high-quality HTS experiments in PubChem covering all important target protein classes for drug discovery. We carefully curated these datasets to have lists of inactive and confirmed active molecules. We make these datasets through PubChem available for benchmarking QSAR models (Butkiewicz et al. 2017, 2013). Data statistics can be seen in Table 1.

**Table 1:** Statistics of datasets used in the experiment. The datasets feature in the large data size, highly imbalanced labels, and diverse protein targets. Datasets are identified by their PubChem Assay ID (AID).

| Protein Target Class | Protein Target (PubChem AID) | Total # of Graphs | # of Active Labels | Per Graph Avg. # of Nodes (Edges) |
| --- | --- | --- | --- | --- |
| GPCR | Orexin1 Receptor (435008) | 218,156 | 233 | 45.14 (94.37) |
|  | M1 Muscarinic Receptor Agonists (1798) | 61,832 | 187 | 43.60 (91.37) |
|  | M1 Muscarinic Receptor Antagonists (435034) | 61,755 | 362 | 43.61 (91.41) |
| Ion Channel | Potassium Ion Channel Kir2.1 (1843) | 301,490 | 172 | 44.41 (92.81) |
|  | KCNQ2 Potassium Channel (2258) | 302,402 | 213 | 44.44 (92.88) |
|  | Cav3 T-type Calcium Channels (463087) | 100,874 | 703 | 43.75 (91.57) |
| Transporter | Choline Transporter (488997) | 302,303 | 252 | 44.46 (92.90) |
| Kinase | Serine/Threonine Kinase 33 (2689) | 319,789 | 172 | 44.85 (93.70) |
| Enzyme | Tyrosyl-DNA Phosphodiesterase (485290) | 341,304 | 281 | 46.13 (96.50) |

### 5.3 Baselines

We benchmark our method, **MolKGNN**, in comparison to five other methods. **SchNet** (Schütt et al. 2017), **SphereNet** (Liu et al. 2021), **DimeNet++** (Gallicchio and Micheli 2020), **ChIRo** (Adams, Pattanaik, and Coley 2021) and **KerGNN** (Feng et al. 2022). The first four are GNNs for molecular representation learning. The last one is a GNN that is architecturally similar to ours. Another similar work (Cosmo et al. 2021) is excluded from benchmarking due to no publicly available code at the time of writing.

### 5.4 Evaluation Metrics

Two metrics are used to evaluate our methods:

- Logarithmic Receiver-Operating-Characteristic Area Under the Curve with the False Positive Rate in [0.001, 0.1] (**logAUC**_[0.001,0.1]_): Ranged logAUC (Mysinger and Shoichet 2010) is used because only a small percentage of molecules predicted with high activity can be selected for experimental tests in consideration of cost in a real-world drug campaign (Butkiewicz et al. 2017). This high decision cutoff corresponds to the left side of the Receiver-Operating-Characteristic (ROC) curve, i.e., those False Positive Rates (FPRs) with small values. Also, because the threshold cannot be predetermined, the area under the curve is used to consolidate all possible thresholds within a certain FPR range. Finally, the logarithm is used to bias towards smaller FPRs. Following prior work (Mendenhall and Meiler 2016; Golkov et al. 2020), we choose to use logAUC_[0.001,0.1]_. A perfect classifier achieves a logAUC_[0.001,0.1]_ of 1, while a random classifier reaches a logAUC_[0.001,0.1]_ of around 0.0215, as shown below:

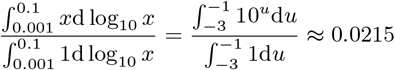
- Receiver-Operating-Characteristic Area Under the Curve (**AUC**): We include AUC since this has historically been used as a general purpose evaluation metric for graph classification (Wu et al. 2018). Comparison with AUC also highlights the fact that overall performance (ranking) of methods according to AUC may not align well with that of the domain specific evaluation metric, i.e., logAUC_[0.001,0.1]_.

### 5.5 Training Details

The datasets are split into 80%/10%/10% for training, validation and testing, respectively. Due to the large size of the datasets and limited computation resources, we reduce our training sets. We use 10,000 randomly selected inactive-labeled samples while keeping all the active-labeled samples for training (following (Mendenhall and Meiler 2016)). The validation and testing sets remain the same and more details of these reduced training sets found in the supplementary material. We leave exploring the full dataset and benchmarking against non-GNN methods as one future direction. To cope with the highly-imbalanced datasets, we use weighted sampling to oversample the active-labeled data in each batch. The codes are provided in the supplementary materials and are implemented using PyTorch (Paszke et al. 2019) and PyG (Fey and Lenssen 2019). More training details can be found in the supplementary material.

### 5.6 Result

From Table 2, we can see MolKGNN achieves superior results in recovering the active molecules with a high decision threshold. This demonstrates the applicability of MolKGNN in a real-world scenario. KerGNN has the worst performance, which aligns with our arguments in Section 2.1 that semantic similarity is more useful than structural in drug discovery. Moreover, we find MolKGNN also performs on par with other GNN in terms of AUC (see Table 3), which demonstrates its potentials applicability beyond drug discovery in a general setting. It is worth noting that different ranking of models are observed in the two tables. This demonstrates that a generally good performing model measured by AUC could potentially perform bad in a specific false positive rate region. Additionally, it highlights the ability of the proposed model to perform well in the application-related metric and indicates its practical significance. Finally the results prompt us to wonder if 3D model can indeed process more information than 2.5D model. The supplementary material contains a discussion on 2.5D vs. 3D models.

**Table 2:** Comparison of logAUC_[0.001,0.1]_ between models. The performance is better when the value is higher. Reported are the mean values over five runs, with standard deviation.

| PubChem AID | MolKGNN (ours) | SchNet | SphereNet | DimeNet++ | ChiRo | KerGNN |
| --- | --- | --- | --- | --- | --- | --- |
| 435008 | $0.255 \pm 0.014$ | $0.187 \pm 0.027$ | $0.215 \pm 0.024$ | $0.203 \pm 0.047$ | $0.168 \pm 0.019$ | $0.147 \pm 0.015$ |
| 1798 | $0.174 \pm 0.029$ | $0.195 \pm 0.025$ | $0.196 \pm 0.035$ | $0.208 \pm 0.035$ | $0.165 \pm 0.040$ | $0.078 \pm 0.042$ |
| 435034 | $0.227 \pm 0.022$ | $0.246 \pm 0.020$ | $0.230 \pm 0.034$ | $0.235 \pm 0.044$ | $0.211 \pm 0.023$ | $0.179 \pm 0.045$ |
| 1843 | $0.362 \pm 0.033$ | $0.358 \pm 0.037$ | $0.258 \pm 0.048$ | $0.284 \pm 0.034$ | $0.326 \pm 0.010$ | $0.292 \pm 0.027$ |
| 2258 | $0.301 \pm 0.028$ | $0.240 \pm 0.037$ | $0.380 \pm 0.037$ | $0.340 \pm 0.032$ | $0.251 \pm 0.010$ | $0.195 \pm 0.020$ |
| 463087 | $0.390 \pm 0.056$ | $0.332 \pm 0.022$ | $0.399 \pm 0.011$ | $0.389 \pm 0.026$ | $0.258 \pm 0.019$ | $0.150 \pm 0.011$ |
| 488997 | $0.303 \pm 0.027$ | $0.319 \pm 0.017$ | $0.309 \pm 0.029$ | $0.315 \pm 0.011$ | $0.193 \pm 0.029$ | $0.081 \pm 0.023$ |
| 2689 | $0.415 \pm 0.020$ | $0.324 \pm 0.020$ | $0.401 \pm 0.016$ | $0.367 \pm 0.049$ | $0.351 \pm 0.048$ | $0.264 \pm 0.017$ |
| 485290 | $0.498 \pm 0.015$ | $0.333 \pm 0.047$ | $0.450 \pm 0.039$ | $0.463 \pm 0.040$ | $0.295 \pm 0.068$ | $0.223 \pm 0.026$ |
| Average | 0.325 | 0.282 | 0.315 | 0.312 | 0.247 | 0.179 |
| Avg. Rank | **2.333** | 3.222 | 2.556 | 2.556 | 4.556 | 5.778 |

**Table 3:** Comparison of AUC between models. The performance is better when the value is higher. Reported are the mean values over five runs, with standard deviation.

| PubChem AID | MolKGNN (ours) | SchNet | SphereNet | DimeNet++ | ChiRo | KerGNN |
| --- | --- | --- | --- | --- | --- | --- |
| 435008 | $0.836 \pm 0.012$ | $0.820 \pm 0.009$ | $0.794 \pm 0.026$ | $0.787 \pm 0.028$ | $0.797 \pm 0.015$ | $0.806 \pm 0.017$ |
| 1798 | $0.721 \pm 0.027$ | $0.707 \pm 0.007$ | $0.655 \pm 0.025$ | $0.649 \pm 0.028$ | $0.683 \pm 0.052$ | $0.663 \pm 0.041$ |
| 435034 | $0.816 \pm 0.028$ | $0.838 \pm 0.009$ | $0.836 \pm 0.014$ | $0.834 \pm 0.019$ | $0.822 \pm 0.017$ | $0.821 \pm 0.016$ |
| 1843 | $0.879 \pm 0.025$ | $0.896 \pm 0.012$ | $0.875 \pm 0.021$ | $0.857 \pm 0.011$ | $0.881 \pm 0.010$ | $0.906 \pm 0.020$ |
| 2258 | $0.806 \pm 0.019$ | $0.792 \pm 0.020$ | $0.801 \pm 0.042$ | $0.821 \pm 0.025$ | $0.782 \pm 0.018$ | $0.766 \pm 0.024$ |
| 463087 | $0.895 \pm 0.003$ | $0.910 \pm 0.005$ | $0.904 \pm 0.005$ | $0.902 \pm 0.009$ | $0.891 \pm 0.004$ | $0.859 \pm 0.009$ |
| 488997 | $0.866 \pm 0.018$ | $0.831 \pm 0.012$ | $0.822 \pm 0.017$ | $0.839 \pm 0.023$ | $0.817 \pm 0.019$ | $0.757 \pm 0.044$ |
| 2689 | $0.906 \pm 0.019$ | $0.905 \pm 0.021$ | $0.867 \pm 0.021$ | $0.832 \pm 0.016$ | $0.919 \pm 0.017$ | $0.912 \pm 0.013$ |
| 485290 | $0.866 \pm 0.012$ | $0.893 \pm 0.011$ | $0.879 \pm 0.021$ | $0.884 \pm 0.016$ | $0.816 \pm 0.015$ | $0.853 \pm 0.009$ |
| Average | 0.843 | 0.844 | 0.826 | 0.823 | 0.823 | 0.816 |
| Avg. Rank | 2.889 | **2.111** | 3.778 | 3.889 | 4.000 | 4.222 |

### 5.7 Investigation of Interpretability

We train a simple autoencoder architecture for interpreting kernels. The encoder is the same as the one used in MolKGNN to convert node features into the node embedding via batch normalization. The decoder converts the node embedding back to the corresponding atomic number. This encoder can be used to translate the node embedding in the kernels into atomic numbers. We examine the learned kernels and below is one example that demonstrates the interpretability of our model. Currently we only examine the first layer and the node attributes of the kernels, but our kernels offer the potentials for retrieving more complicated pattern and we leave the investigation of that for future works.

In Figure 5, the learned pattern shows a center atom of carbon surrounded by neighboring three fluorine and another carbon. An examination of the training set reveals several molecules displaying this pattern. This finding corresponds to the domain knowledge: the pattern is known as the trifluoromethyl group in medicinal chemistry and has been used in several drugs (Yale 1958).

**Figure 5:**
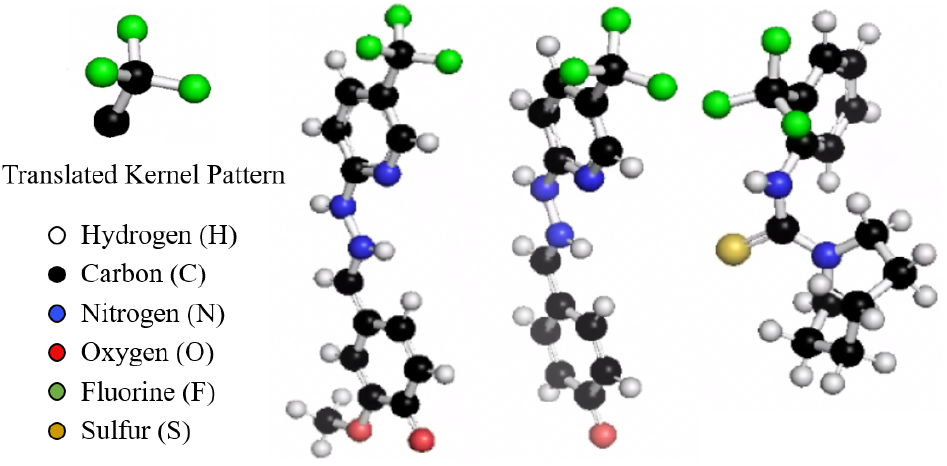
Visualization of a learned kernel of MolKGNN when trained on AID 2689. This first highlights the interpretability of our model, while also providing an example of a learned kernel aligning with domain knowledge/expectations, since this is a known important substructure in medicinal chemistry, i.e., the trifluoromethyl group (shown in three example molecules).

### 5.8 Ability to Distinguish Chirality

We further experiment on the expressiveness of our model to determine whether it is able to distinguish chiral molecules. We use the CHIRAL1 dataset (Pattanaik et al. 2020) that contains 102,389 enantiomer pairs for a single 1,3-dicyclohexylpropane skeletal scaffold with one chiral center. The data is labeled as R or S stereocenter and we use accuracy to evaluate the performance. For comparison, we use GCN (Kipf and Welling 2016) and a modified version of our model, MolKGNN-NoChi, that removes the chirality calculation module. Our experiments observed GCN and MolKGNN-NoChi achieve 50% accuracy while MolKGNN achieves nearly 100%, which empirically demonstrates our proposed method’s ability to distinguish chiral molecules.

### 5.9 Ablation Study

#### Component of *ϕ*(*S, S*’)

We conduct an ablation study on the three compnents of *ϕ*(*S, S*’), i.e., *ϕ*_cs_, *ϕ*_ns_, *ϕ*_es_. From the result in Figure 6, we observe that the removal of any of the components has a negative impact on logAUC_[0.001,0.1]_. In fact, the impact is bigger for logAUC_[0.001,0.1]_ than AUC in terms of the percentage of performance change. Note that in some cases such as the removal of *ϕ*_es_, there is an increase in performance according to AUC, but this would significantly hinder the logAUC_[0.001,0.1]_ metric.

**Figure 6:**
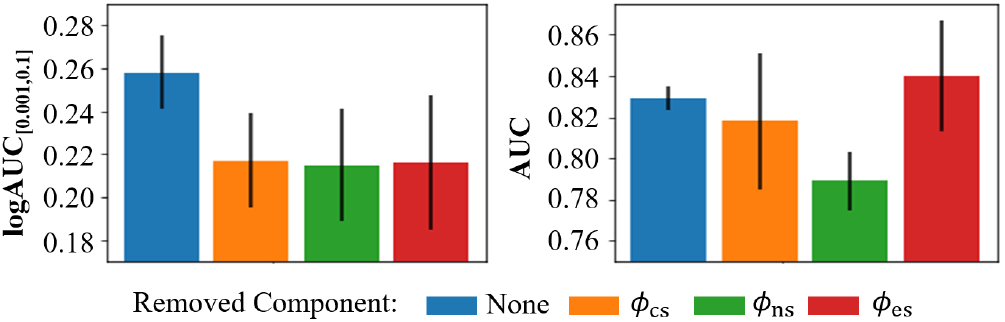
Ablation study result for *ϕ*(*S, S*’) components using AID 435038. Reported are average values over three runs, with standard deviation.

#### Kernel Number

In addition, an ablation study is conducted to study the impact of kernel numbers (results in Figure 7). When the number of kernels is too small (< 5), it greatly impacts the performance. However, once it is large enough to a certain point, a larger number of kernels has little impact on the performance.

**Figure 7:**
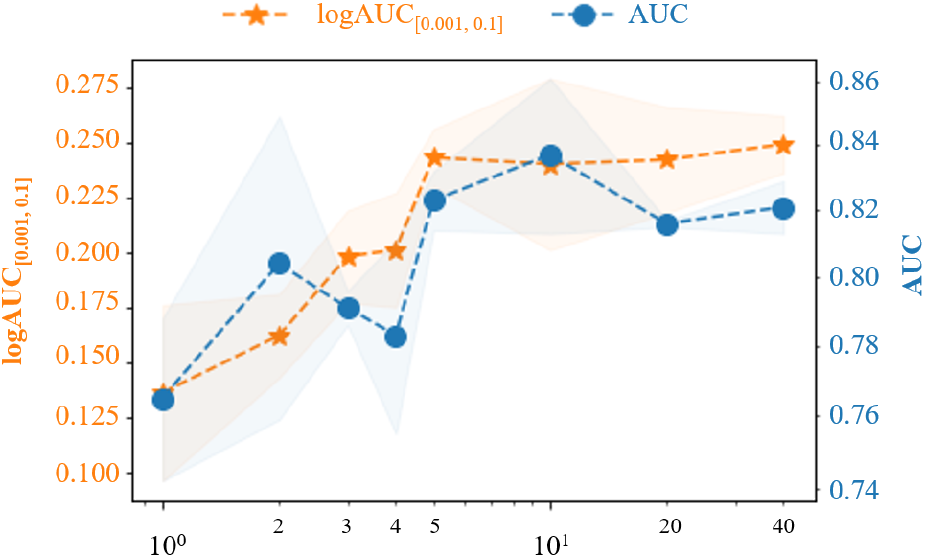
Performance for different kernel numbers using AID 435008. The number shown is applied to kernels of all degrees. Results are average values over three runs, with standard deviation.

## 6 Discussion

### 6.1 Computation Complexity

It may seem to be formidable to enumerate all possible matchings described in Section 4.1. However, most nodes only have one neighbor (e.g., hydrogen, fluorine, chlorine, bromine and iodine). Take AID 1798 for example, 49.03%, 6.12%, 31.08% and 13.77% nodes are with one, two, three and four neighbors among all nodes, respectively. For nodes with four neighbors, only 12 out of 24 matchings need to be enumerated because of chirality (Pattanaik et al. 2020).

Since the adjacency matrix of molecular graphs are sparse, most GNNs incur a time complexity of 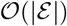. And as analyzed above, the permutation is bounded by up to four neighbors (12 matchings). Thus, finding the optimal matching has a time complexity of 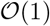. The calculation of molecular convolution is linear to the number of *K* kernels and hence has a time complexity of 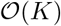. Overall, our method takes a computation time of 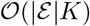.

## 7 Conclusion

In this work, we introduce a new GNN model named MolKGNN to address the QSAR modeling problem. MolKGNN utilizes a newly-designed molecular convolution, where a molecular neighborhood is compared with a molecular kernel to output a similarity score, which is used as the new atom embedding for the next layer. Comprehensive benchmarking is conducted to evaluate MolKGNN. Well-curated datasets that consist of experimental HTS data from diverse protein target classes are used for the evaluation. The datasets are highly imbalanced, which highlights the scarcity of positive signals in this real-world problem. For evaluation, not only do we use traditional AUC, but also logAUC_[0.001,0.1]_, to evaluate the method’s performance under a high cutoff condition. This high-cutoff condition is typical of real-world applications and demonstrates the applicability of MolKGNN to drug discovery. Moreover, this paper provides a theoretical justification and experimental demonstration that MolKGNN is able to distinguish chiral molecules while providing interpretability for its results. As a future direction we aim to leverage self-supervised learning on graphs (Jin et al. 2020; Wang, Jin, and Derr 2022) to further improve our prediction results.

## 8 Supplementary Material

### 8.1 Experiment Details

#### Data Preprocessing

We preprocessed the input SMILES strings to Structure-Data Files (SDFs). The preprocessed SDFs have been made publicly available and can be found at: https://doi.org/10.6084/m9.figshare.20539893.v1. The dataset is specified by its PubChem Asssay ID (AID) (Wang et al. 2012). Prepossessing to the original data includes converting SMILES strings to 3D SDF files, generating 3D conformation, and filtering. Conversion from SMILES to SDF files is done using Open Babel (O’Boyle et al. 2011), version 2.4.1. Conformations are generated using Corina (Gasteiger, Rudolph, and Sadowski 1990), version 4.3. Molecules are further filtered with validity, duplicates with BioChemical Library (BCL) (Brown et al. 2022).

#### Data Split

The datasets are randomly split into 80%/10%/10% for training, validation, and testing respectively. We then shrink the training set to contain only 10,000 inactive-labeled molecules, while keeping all active-labeled molecules. This shrinking technique was previously used by (Mendenhall and Meiler 2016) By shrinking the training data size, we can shorten the training time given the limited computational resources, while keeping most active signal because only those predicted active molecules will be experimentally validated and hence we focus on those. We did an empirical study on the shrinking effect on AID 2258 (302,402 molecules). Results are in Table 4. We can see there is indeed a decrease of performance in terms of logAUC_[0.001,0.1]_. We leave the benchmarking of the full dataset in a future study.

**Table 4:** Results comparison between shrinked and full training set using MolKGNN over three runs for AID 2258.

| Inactive Training Size | $\log\text{AUC}_{[0.001,0.1]}$ | AUC |
| --- | --- | --- |
| 10K Sample | $0.296 \pm 0.026$ | $0.820 \pm 0.021$ |
| Full | $0.384 \pm 0.003$ | $0.816 \pm 0.030$ |

#### Training Details

To overcome the highly-imbalanced problem, we sample the training data in each batch according to the inverse frequency of the label occurence in the training set. For example, if the active label appear at 1% rate in the training set, it has a sampling weight of 1/0.01 =100, while if the inactive label appear 99% of time in the training set, it gets a sampling weight of 1/0.99 ≈ 1.01. The active-labeled data are thus roughly 100 times more likely to be sampled than inactive-labeled data in each batch. A linear layer is used for the *f* (·) in Equation 11.

### 8.2 Hyperparameters

Table 5 contains details for the hyperparameter search space for MolKGNN and additional training details are in Table 6. For other benchmarking models except KerGNN, we use the same hyperparameters from their codes. For KerGNN, we observed that using the default hyperparameter setting achieves significantly low performance on our well-curated datasets and hence we further tune its hyperparamters as follows: batch size {64,128}, the hidden layer units {16, 32}.

**Table 5:** Hyperparameter search space used for MolKGNN.

| Hyperparameter | Search Space |
| --- | --- |
| Hidden Dimension | {32, 64} |
| Batch Size | {16, 32} |
| # Layers | {1, 2, 3, 4, 5} |
| Peak Learning Rate | {5e-1, 5e-2, 5e-3, 5e-4} |
| Dropout | {0.1, 0.2, 0.3} |

**Table 6:** Hyperparameters used for MolKGNN.

| Hyperparameter | Value |
| --- | --- |
| Node Feature Dimension | 28 |
| Edge Feature Dimension | 7 |
| Hidden Dimension | 32 |
| Batch Size | 16 |
| # Layers | 4 |
| # of Kernels of Degree 1 | 10 |
| # of Kernels of Degree 2 | 20 |
| # of Kernels of Degree 3 | 30 |
| # of Kernels of Degree 4 | 50 |
| Warmup Steps | 300 |
| Peak Learning Rate | 5e-3 |
| End Learning Rate | 1e-10 |
| Weight Decay | 0.001 |
| Epochs | 20 |
| Dropout | 0.2 |

### 8.3 Featurization

Different models have different ways of featurization. We use the original features reported in the original papers for each model used in the benchmarking. Our featurization is adapted from (Coley et al. 2017). Rdkit(version 2022.3.4) (Landrum et al. 2013) is used for the featurization. See Table 7 and 8 for details.

**Table 7:** Node features **X**_*υ*_, for *υ*.

| Indices | Description |
| --- | --- |
| 0-11 | One-hot encoding of element type: H, C, N, O, F, Si, P, S, Cl, Br, I, other |
| 12-15 | One-hot encoding of node degree: 1, 2, 3, 4 |
| 16 | Formal charge |
| 17 | Is in a range |
| 18 | Is aromatic |
| 19 | Explicit valence |
| 20 | Atom mass |
| 21 | Gasteiger charge |
| 22 | Gasteiger H charge |
| 23 | Crippen contribution to logP |
| 24 | Crippen contribution to molar refractivity |
| 25 | Total polar surface area contribution |
| 26 | Labute approximate surface area contribution |
| 27 | EState index |

**Table 8:** Edge features **E**_*υu*_ for *e*_*υu*_.

| Indices | Description |
| --- | --- |
| 0 | Is aromatic |
| 1 | Is conjugate |
| 2 | Is in a range |
| 3-6 | One-hot encoding of bond type: 1, 1.5, 2, 3 |

### 8.4 2.5D vs 3D

While many previous work have attempted to develop 3D models by including distance, angles, torsions into their model designs (Schütt et al. 2017; Klicpera, Groß, and Günnemann 2020; Liu et al. 2021), we demonstrate that 2.5D model can achieve comparable results in terms of AUC, or even better results in terms of logAUC[0.001,0.1]. We provide the explanation of why a model with seemly less information can accomplish this from a chemistry perspective: The bond lengths/angles have little variations given the certain involving atom identities and bond types (Gordy 1947; Gillespie 1960). Moreover, different than determining bond lengths/angles experimentally, many programs such as Corina (Gasteiger, Rudolph, and Sadowski 1990) that converts SMILES to 3D SDF using standard bond lengths^2^, which stay the same in different molecules. Hence bond lengths/angles provide little additional information in distinguishing different molecules. This can also been seen by the fact that an experienced chemist can just look at a molecular structure and know certain properties of the molecule, without the need to know the exact bond lengths/angles. Nevertheless since our model has the potential to integrate bond length and angles in the *ϕ*(*S, S*’), we plan to include those for comparison in the future studies.

On the other hand, molecules can have different conformations as a result of the single bond rotation. The same molecule with different conformation consequently has different sets of torsions. However, the pharmacological activity is usually linked with few conformations (binding conformations) and hence related to certain sets torsions. It seems that knowing torsion could potentially help the activity prediction. Nevertheless, knowing which conformation is the binding conformation is a challenging task. A set of torsions related with a wrong predicted binding conformation is detrimental to the model performance. Hence we decide to build a conformation-invariant model and exclude torsion to circumvent this problem.

## Footnotes

1 The lastest one is from 2021.

2 This is explicitly mentioned in its manual: https://mnam.com/wp-content/uploads/2021/10/corina_classic_manual.pdf

